# Label-free analysis of bacterial growth and lysis at the single-cell level using droplet microfluidics and object detection-oriented deep learning

**DOI:** 10.1101/2023.06.27.546533

**Authors:** Anuj Tiwari, Nela Nikolic, Vasileios Anagnostidis, Fabrice Gielen

## Abstract

Bacteria identification and counting at the small population scale is important to many applications in the food safety industry, the diagnostics of infectious diseases and the study and discovery of novel antimicrobial compounds. There is still a lack of easy to implement, fast and accurate methods to count populations of motile cells at the single-cell level. Here, we report a label-free method to count and localize bacterial cells freely swimming in microfluidic anchored picolitre droplets. We used the object detection oriented YOLOv4 deep learning framework for cell detection from bright-field images obtained with an automated Z-stack setup. The neural network was trained to recognize *Escherichia coli* cell morphology with an average precision of approximately 84%. This allowed us to accurately identify individual cell division events, enabling the study of stochastic bacterial growth starting from initial populations as low as one cell. This work also demonstrates the ability to study single cell lysis in the presence of T7 lytic bacterial viruses (phages). The high precision in cell numbers facilitated the visualization of bacteria-phage interactions over timescale of hours, paving the way towards deciphering phage life cycles in confined environments.

## Introduction

The early detection of bacterial presence is of primordial importance for human and animal welfare [1, 2]. In particular, the presence of food microbial contaminants, or antimicrobial resistant strains in hospital settings constitutes a severe threat to human health with up to 15% of all hospitalized patients affected by healthcare-associated infections [3-5]. Rapid identification strategies and their translation into diagnostic devices hold the promise to improve prevention and alleviate healthcare burden [6]. In addition, early elimination of pathogenic bacteria would reduce risks that they develop defence mechanisms under long-term selection pressure (e.g. from a specific food source, antibiotic regimen) [7, 8].

With rising numbers of antimicrobial resistant strains, it is also important to rapidly identify and assess the efficacy of bacteriolytic agents without the need for long culturing times (which may not be feasible for non-culturable strains) [9, 10].

Amongst alternatives to antibiotics to combat bacterial infections, bacterial viruses or phages lyze bacteria in a highly-specific manner, preserving healthy microbiomes while making safe and effective patient-specific treatment a possibility [11, 12]. Compassionate use has already seen some significant success in treating recurrent infections [13, 14]. However, current gold standard assays to quantify phage efficacy such as the plaque assay are slow and labour intensive [15]. Methods that can rapidly assess the lytic potential of phages without requiring extensive bacterial culture are still needed. Likewise, platforms that inform on possible bacterial resistance to phages will play an important role to understand future clinical use for phage therapy [16].

Testing antimicrobials on a small subset of cells would accelerate time to results which could shorten the time to deliver adequate treatment. Current methods to study bacterial populations down to the single-cell level focus on either growth of bacterial cells in 2D on the surface of agar plates, confined within microchannels or in small liquid cultures [17-21]. 2D cultures produce cell monolayers in early growth stages which are highly suited for high-resolution imaging with the ability to track cell divisions and individual lineages for an in-depth understanding of growth kinetics. This has allowed the development of analytical models for predicting bacterial growth from a single or a few cells [22, 23].

In their natural environment, bacteria may be found in water-filled microcavities or in condensation droplets in which they can proliferate without attachment to a solid substrate. For some strains, the ability to swim is an essential phenotypic trait which help them colonize new habitats or respond to physical or chemical cues [24]. Understanding unconstrained bacterial growth in 3D is therefore desirable to quantify cell lifestyle which includes non-surface attached aggregates or biofilms [25]. Growing cells within microfluidic or sessile droplets enable long-term visualization of a small group of cells confined within picolitre volumes [19]. Many methods have been developed to ensure long-term, stable imaging, for example by anchoring droplets in multilayer devices, or using geometrical traps [26-29]. In addition, numerous methods have been developed for counting cells using various imaging modalities in 2D formats [30, 31]. In the context of 3D motion, there is an unavoidable trade-off between technical simplicity, counting accuracy and time-to-result [32].

The use of fluorescence markers to count and track individual cells result in high sensitivity and signal-to-noise ratio, and has been historically successful for assessing bacterial population dynamics [33]. On the other hand, fluorescently labelling bacteria may interfere with the cell biology and would increase the number of steps of the overall sample preparation. For these reasons, it is desirable to use label-free imaging (e.g., transmitted light, phase contrast, dark field) on unaltered cell samples, simplifying overall procedures and making them generally more applicable. Various complex optical systems have been devised to perform 3D object tracking. Multifocal imaging can provide volumetric imaging data and be used in phase contrast or darkfield modes [34, 35]. Digital holographic microscopy with multiple light sources has been used to track freely diffusing *E. coli* bacteria [36]. In phase contrast mode, diffraction patterns can be used to infer localization of bacterial cells [37]. Dark-field imaging with an 87-channel multispectral system has been used to identify several bacterial species [38]. Standard bright-field imaging, although one of the simplest and most widely used modality, typically affords lower contrast-to-noise ratio and does not provide clear volumetric data. Accurate counting of individual, unstained cells over time remains a significant challenge [39]. Even when using low height microchambers relative to cell size, cell identification is difficult as cells can move in and out from the focal plane and orient themselves randomly resulting in various appearances for the same cell. This explains why studies, even when using fluorescent strains, have focussed on obtaining estimates of absolute cell counts for large populations of up to ⁓10,000 cells [20],[21]. Although such approaches provide information on large population growth kinetics, they are not so informative on early-stage population development. In particular, early cell division events may be missed even though they are crucial for informing on cell viability, capturability and adaptation phase.

Cell detection has recently benefited from significant advances made in deep learning algorithms, including from images obtained using bright-field microscopy. Deep learning algorithms such as two-stage (e.g. R-CNN, SPP-Net) and single-stage detectors (e.g. YOLO, SSD) have proven abilities to accurately identify, classify and locate objects in images using manually curated training datasets [40-42]. Efforts towards dissemination of such deep learning models (e.g. through open-access training datasets) have made these methods more widely accessible [39].

In this paper, we demonstrate label-free, accurate counting of freely-swimming bacterial cell populations starting from as few as one cell. We combine standard bright-field imaging with a rapid and automated Z-scanning method which enables detection of 3D positions for cells growing in 8 μm tall, anchored droplets. We use the YOLOv4 object detector to count in-focus cells and obtain accurate cell numbers over time. Finally, we demonstrate the ability of the platform to detect single cell lysis events induced by lytic phages.

## 2. Methods

### 2.1 Bacterial strain and phage lysate preparation

Bacterial strain *Escherichia coli* BW25113 was used for all the experiments in this study. Streaks consisting of single colonies on Lysogeny Broth (LB) agar plates were obtained from a glycerol stock of the strain stored at -80°C. Individual colonies were picked for each experiment and added to a sterile culture tube containing 4 mL of LB media (10 g/L tryptone, 5 g/L yeast extract, 10 g/L NaCl) and incubated overnight at 37°C with shaking at 230 rpm. 40 μL of the overnight culture was added to 4 mL of fresh LB media and cultivated at 37°C and 230 rpm until it reached a desirable OD_600_ reading (optical density measured at 600 nm, serving as a proxy for bacterial population size). Standard colony forming unit (CFU) assays were also carried out for each experiment (Supplementary, section 2.1). Phage T7 lysates were preserved in SM buffer (0.1 M NaCl, 8 mM MgSO4·7H2O, 50 mM Tris-Cl pH 7.5, 0.01% gelatin) at 4°C, with phage titer of 10^9^ PFU/mL (PFU, plaque-forming unit) (Supplementary, section 2.2 and Figure S1). All experiments were performed at room temperature (measured to be around 25 °C) and at 37°C respectively, as *E. coli* is capable of growing in a wide temperature range (8°C-48°C) [43].

For growth experiments, 1 mL aliquots of desired OD_600_ culture were directly utilized for experiments. For lysis experiments, aliquots of OD_600_ cultures were mixed with phage titers in different volumes to obtain a desired multiplicity of infection (MOI). MOI is the ratio between the number of phage particles and number of bacterial cells. The experimental conditions are listed in Table 1 and Table 2.

**Table 1:** Experimental conditions for growth experiments.

| Experiment | OD<sub>600</sub> | CFU density (cells/mL) | Temperature | Initial cell count in droplet |
| --- | --- | --- | --- | --- |
| G1 | 0.19 | 0.6 x 10<sup>8</sup> | Room Temperature | 2 |
| G2 | 0.35 | 2.4 x 10<sup>8</sup> | Room Temperature | 1 |
| G3 | 1.00 | 6.5 x 10<sup>8</sup> | Room Temperature | 5 |
| G4 | 0.29 | 1.8 x 10<sup>8</sup> | 37°C | 4 |
| G5 | 0.30 | 2.1 x 10<sup>8</sup> | 37°C | 3 |
| G6 | 0.35 | 2.2 x 10<sup>8</sup> | 37°C | 2 |

**Table 2:**
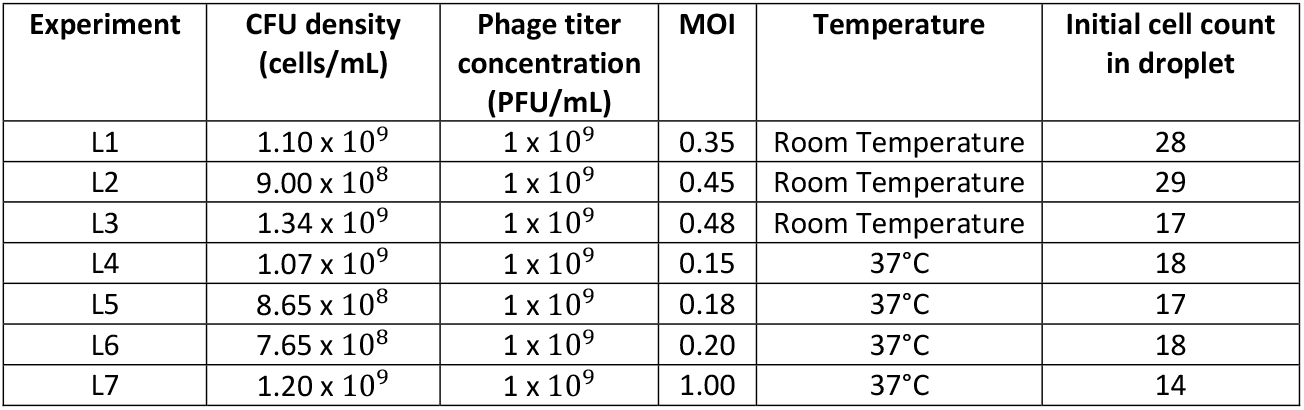
Experimental conditions for lysis experiments.

### 2.2 Microfluidic device fabrication

The two-layer microfluidic device was fabricated using soft lithography with a high-resolution acetate mask (Microlithography Services). Negative photoresist SU8 TF6000 (MicroChem) was patterned on a silicon wafer by exposure to UV light through a transparent film mask. The device consists of two layers with pillars (first layer, 4 μm) and microdroplet traps (second layer, 4 μm). The droplet trap diameter was 60 µm. A detailed procedure is described in a previous work [44]. PDMS (polydimethylsiloxane, Ellsworth) and curing agent were mixed in a ratio of 10:1, poured on the patterned silicon wafer and degassed. To minimize evaporation of droplets during the time-lapse experiments, a small piece of coverslip (thickness 0.15 mm) was placed over the trap array prior to the curing process. The wafer was then cured at 70 °C for 120 minutes. The cured PDMS was cut out and 1 mm holes were punched (Kai Medical) to create inlets for the oil and bacteria culture media. Plasma treatment (Diener Zepto) was used to bond the PDMS to a thin coverslip (22 × 50 mm, 0.13–0.17 mm thick). Finally a solution of 1% v/v silane dissolved in HFE-7500 oil was flushed through the device which was incubated at 70°C for at least 30 minutes.

### 2.3 Droplet generation and trapping

The microfluidic device was placed on the microscope stage and clamped using scotch tape. Droplets were generated on-chip by phase change in which the aqueous phase containing the bacterial cells/bacteria-phage mix was replaced by the oil phase, forming anchored droplets where traps were designed. The continuous phase consisted of 1% (w/v) 008-Fluorosurfactant (RAN Biotechnologies) in HFE-7500 (Fluorochem) oil. The aqueous phase consisted of *E. coli* cultured in growth medium until the indicated OD_600_ (control experiments) or *E. coli*-T7 phage mixture at a specific MOI. A schematic of the microscope setup is shown in Figure 1A. To start the experiments, the aqueous and carrier oil phase were loaded into PTFE tubings (SLS) connected to 1 mL plastic syringes (BD Plastipak) and syringes were mounted on syringe pumps (Nemesys, Cetoni). 100 µL of bacterial cell culture/bacteria-phage mix was aspirated in one of the syringes as aqueous phase. The device was filled with the oil solution to remove all the air inside the trapping chambers. The oil flow was then stopped, and bacteria solution/bacteria-phage mix was flown until the whole trapping array was filled. In turn, the bacteria solution/bacteria-phage mix was stopped, and the oil solution was introduced again to flush the cell sample. This procedure created droplets of the cell sample immobilised in the circular traps as seen in Figure 1B.

**Figure 1:**
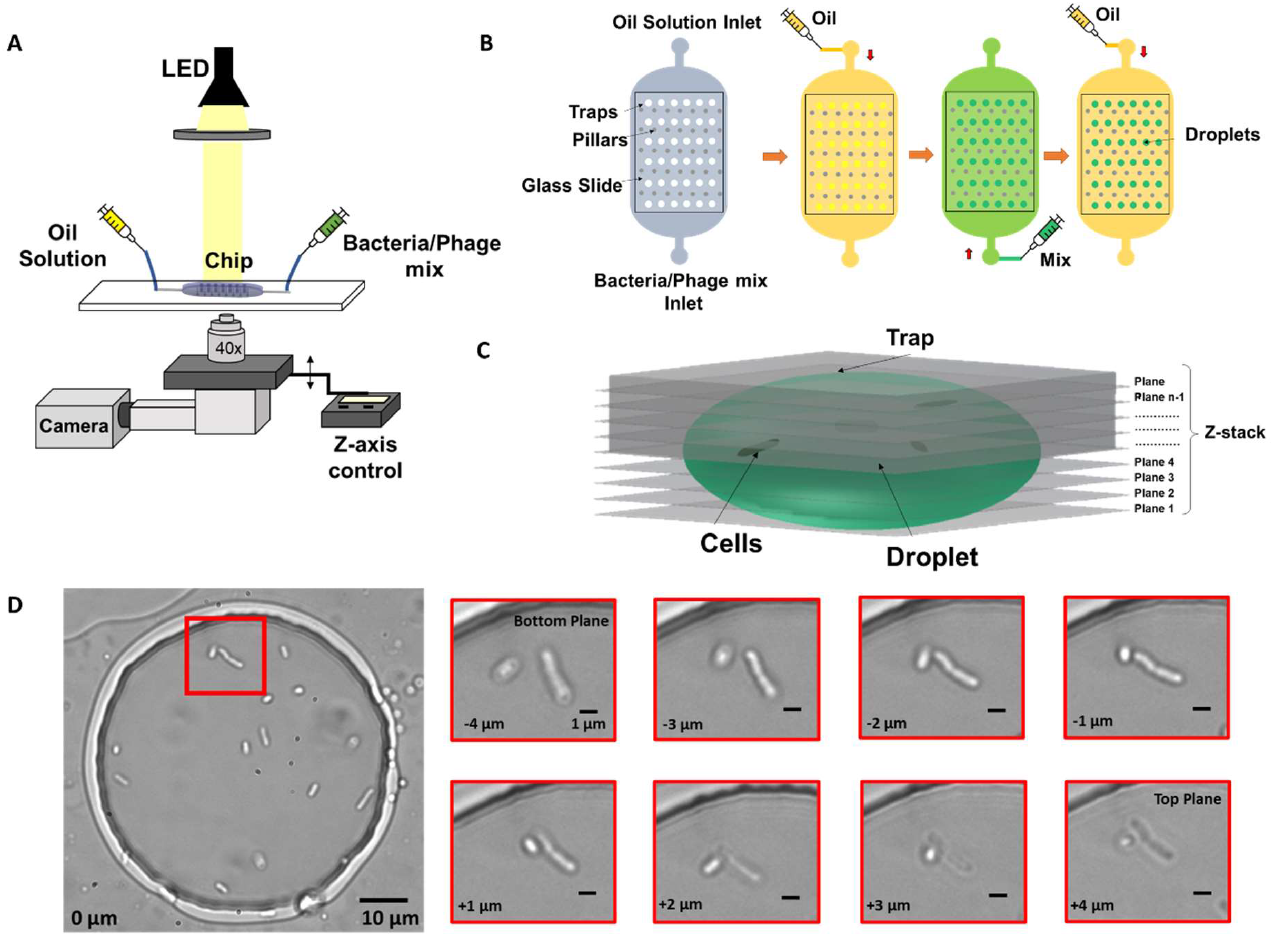
Imaging and microfluidics setup to observe individual bacterial cells trapped in droplets. **(A)** Droplets were illuminated by a collimated white LED light source and imaged in bright-field, using a 40x objective on a microscope with motorized Z-focus control. **(B)** Droplets were generated using a water-in-oil self-digitization method. The microfluidic chip was first primed with oil to remove any air. Bacteria solution or bacteria-phage mix was flushed to replace the oil solution. The oil solution was flushed once more to form droplets in individual traps. **(C)** 3D schematic of Z-stack imaging on a trapped droplet. **(D)** Example image of multiple cells encapsulated in an anchored droplet and zoomed-in image panel showing two cells obtained at eight different Z focal planes. The relative vertical distance between images is indicated at the bottom left of every image.

### 2.4 Imaging

A major challenge for observing individual bacteria by microscopy in 3D environments include their ability to swim away from optimum focus between different planes as well as lateral movement within the said environment. To compensate for this motion and obtain accurate counting, we employed a Z-stack method together with image-based drift correction to image the bacterial cells inside the traps as shown in Figure 1C over timescales of hours. Figure 1D shows an example droplet containing multiple cells and zoomed-in images of cells obtained at different Z focal planes.

An inverted Olympus IX73 semi-motorized microscope was used to image the bacteria trapped in the chambers. The microscope was equipped with a white LED (CoolLED pE-100) and a direct coupling motorized Z-focus with a focus controller (ProScan III, Prior Instruments). We used a 40x objective with a numerical aperture of 0.45 (UPLFLN40X-2, Olympus). The microscope was placed on a vibration damping platform (Newport VIP320X1218-50140). A USB camera (DMK 37AUX287, The Imaging Source) was used to obtain images of droplets of size 720 x 540 pixels encoded on 8 bits.

A Python script was written to control the imaging sequence as well as the movement of the vertical position of the microscope objective Z-axis. Drift correction was utilized to compensate for the possible impact of any external factors such as movement of the clamped device or vibration of the table on which the microscope is set. The Python script enabled us to control the number of stacks, the time gap between the stacks, the height of each step in the stack as well as correct any drift in focus that might occur due to factors mentioned above. The first step in drift correction was to generate an average image calculated from 10 different images of the same chamber. The average image was generated by taking an average of light intensity over 10 images. This average image was then used as a reference for actual stack imaging. Each time a new stack was acquired, the first image of the stack was checked against the average image obtained initially (sum of all absolute values of pixel-to-pixel subtraction). If this global difference between the images was found to be over a certain threshold, the Z-axis would move in either direction (upwards or downwards) until the difference between the new stack image and the average image changed to less than the threshold. If Z-axis movement failed to find suitable correction with vertical movement, a new average image was generated replacing the old image. This process continued over the duration of the experiment. All images within each Z-stack were taken within 2.1 seconds (at the speed limit of the setup) to minimize the effect of cell motion during acquisition. All imaging parameters used in this study can be found in Supplementary section 1.

### 2.5 Deep learning using YOLOv4 for *E. coli* single-cell detection: labelling, training, and detection

YOLOv4 is a one-stage deep learning based object detection framework capable of detecting different classes of objects in images with high speed and accuracy [45]. Images were obtained using the setup mentioned in section 2.4 to create a dataset containing examples of *E. coli* cells in droplets. The dataset consisted of 200 images chosen across different microscopy conditions (light intensity, collimator position) to improve the robustness and versatility of the model. A total of 1670 in-focus cells were manually labelled by drawing bounding boxes. Examples of cells considered in focus can be seen in Figure S2. The labelled data were split into a ratio of 70:30 for training and validation, respectively. The model was trained for 6000 iterations on YOLOv4 darknet using the Google CoLab platform. Trained YOLOv4 weights that exhibited high average precision (AP) and low loss were obtained and converted into TensorFlow format for post processing (c.f. Supplementary sections 4 and 5). The converted YOLOv4 model was used to perform detections on all experimental data. For every raw image input, the model outputs a detection image containing bounding boxes around cells and a NumPy file containing the information on the coordinates of these bounding boxes.

### 2.6 Utilising detections for data analysis

A counting method extracting the number of *E. coli* cells per droplet using Z-stack information was implemented using MATLAB. Every image part of the Z-stack had associated YOLOv4 detections. In a typical experiment, a selected droplet was imaged across more than 20 focal planes, each spaced by 0.5 microns. The plane with the maximum number of cell detections was used to obtain an initial cell count. However, the same cell may be detected in multiple planes due to its changing orientation and motility. It took 75 milliseconds to image two consecutive slices that were 0.5 microns apart. We experimentally determined that a cell would have moved on average less than 4 microns in the same timeframe (c.f. analysis in Figure S5). If, for a slice different than the initial one, a cell was detected at a position sufficiently close to a cell already counted, we assumed that it was the same cell and therefore was registered only once. However, if two cells lied in the same plane, the algorithm would count both, independent of their relative location. After all slices were processed, a final cell count and location was obtained (referred to as “detection count”). This process was repeated for all time points, enabling tracking of cell counts in a given droplet.

The raw “Detection count” data was filtered to extract accurate times for individual cell division events. We implemented a function that decreased detection noise and rounded the cell counts to the local maximum values. Briefly, we applied a moving average function for every 3 data points to attenuate sudden spikes. Inaccuracies in cell detection count can occur, mainly because of the positioning of the cells with respect to the droplet interface. Figure S7 displays examples of false detections. This results in local fluctuations across time series. Consequently, a single change in the detection count may not be reflective of an actual change in cell number. Therefore, we only updated the cell count whenever the same detection number was repeated N times within a given time interval (referred to as the N^th^ maximum method). This filtered cell count was termed “processed count” in Figures 2C and Figure 3.

**Figure 2:**
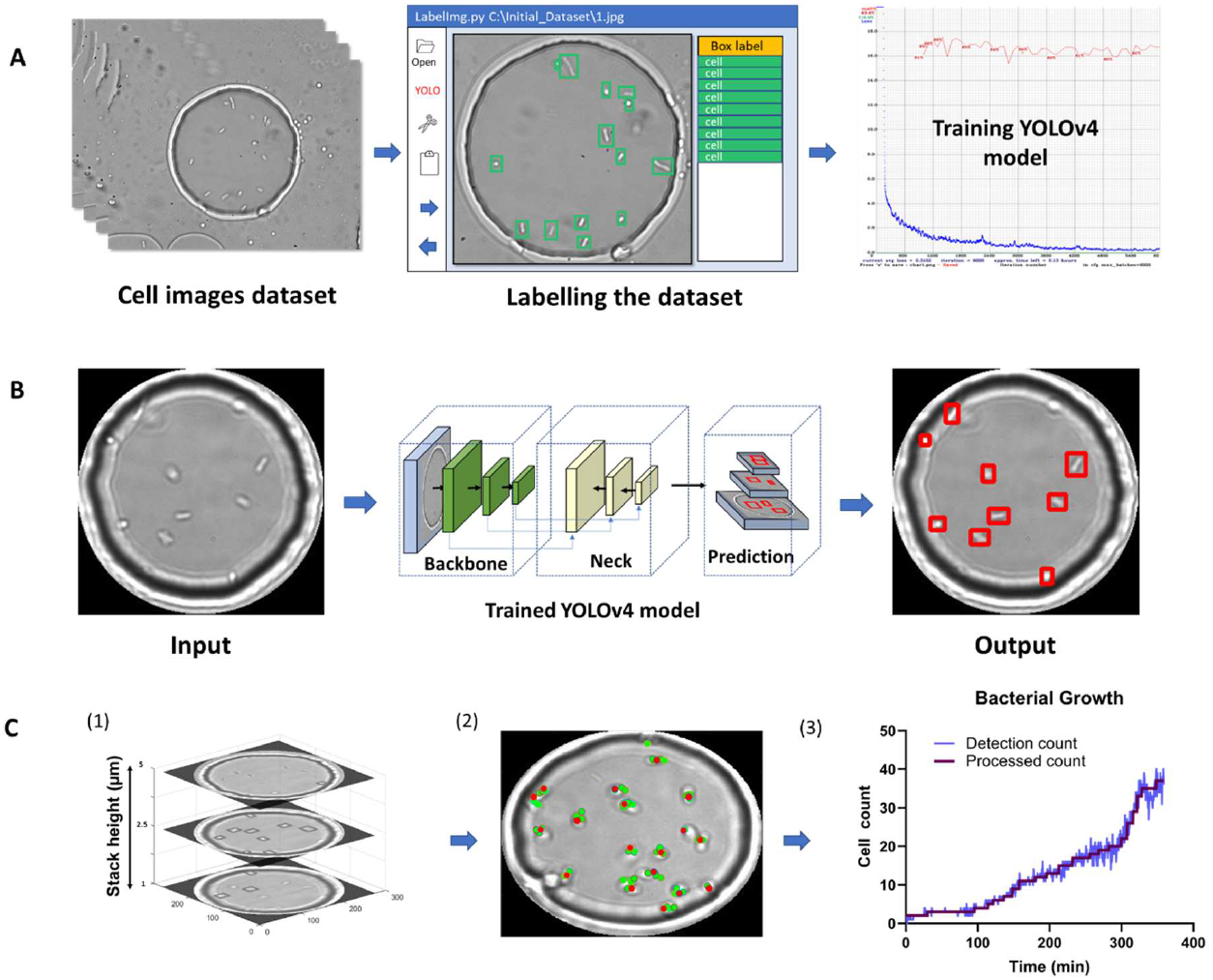
YOLOv4 model training and data analysis. **(A)** Flow chart of labelling procedure involving dataset acquisition, labelling using LabelImg and training using YOLOv4. **(B)** Flow chart showing image masking, image passing through YOLOv4 and cell detection output image. **(C)** MATLAB analysis flowchart. (1) Example Z-stack with cells detected for 3 planes. (2) The location of detected single cells overlaid on one of the Z-stack images are represented by red dots (Supplementary Figure S5). (3) Cell counts over time. The figure was plotted using the process described in (2) for every time point. The final processed count was generated using the 3^rd^ maximum method (see methods).

**Figure 3:**
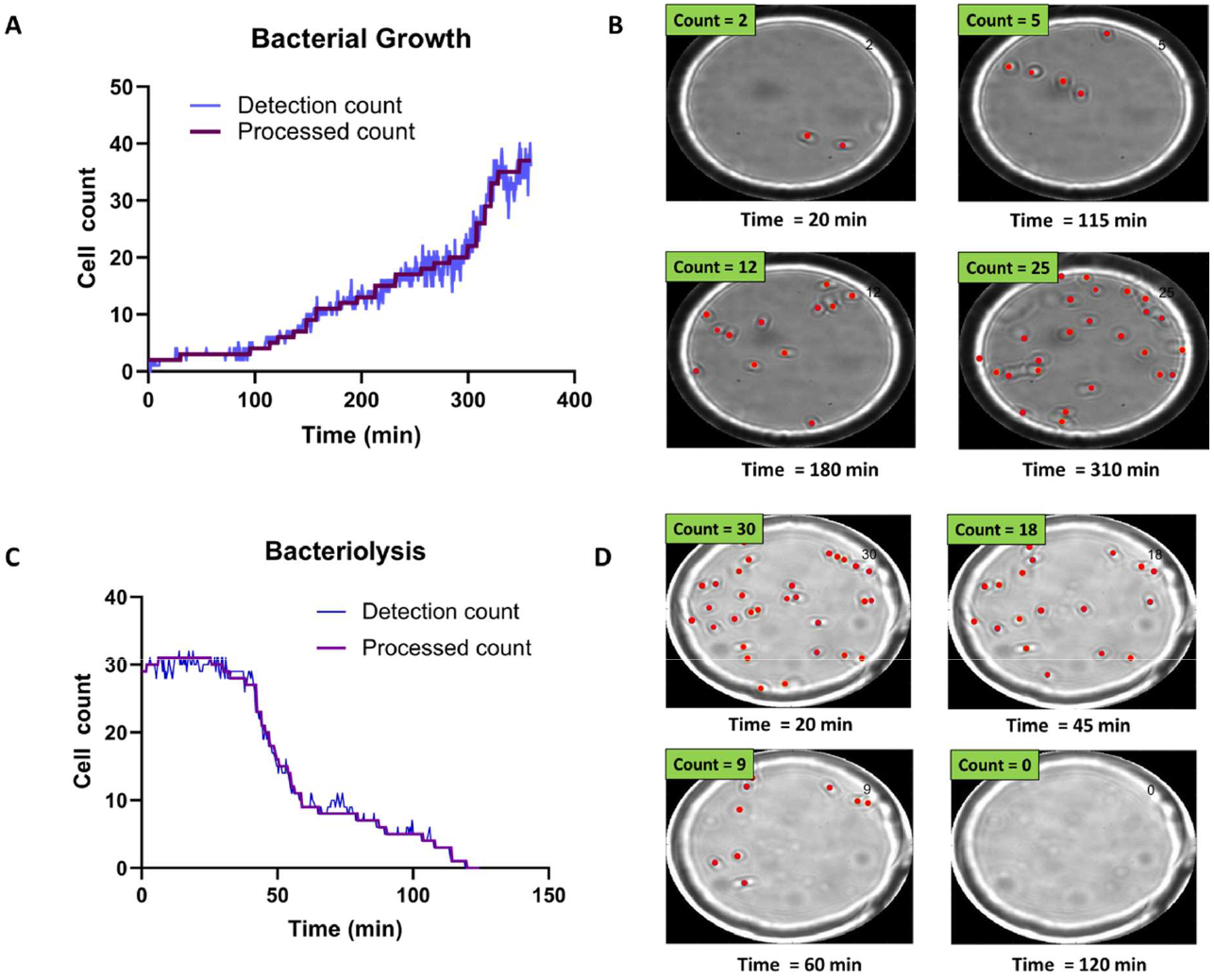
Time-resolved bacterial growth and phage-induced bacteriolysis. **(A)** Example of detection and processed counts for *E. coli* population growth. Starting from a single *E. coli* cell, we tracked the growth of the population. **(B)** Selected images showing corresponding detections at different time points (Movie S1) **(C)** Example of detection and processed counts for lysis of individual *E. coli* cells. **(D)** Selected images showing corresponding detections at different time points (Movie S2).

Similarly, for lysis experiments, if a decrease in cell count was observed N times, the processed cell count was decreased to the lowest count value (‘N^th^ minimum method’). This allowed us to generate the step count increases/decreases and obtain accurate cell division and lysis times for all growth and lysis experiments.

Figure 2 summarizes the image processing workflow, including YOLOv4 training, detections across slices and post-processing to obtain final cell counts and locations.

### 2.7 Calculation of doubling times using processed bacterial counts

Doubling times for bacterial population were calculated using the slope of the bacterial count plots. We manually identified the portion of the plots with highest slope and excluded at least the first 20 and 70 minutes for the 37 °C and room temperature experiments respectively (individual fits are shown in Figure S10). The resulting linear fits helped us obtain the specific growth rates and doubling times using the following equation:

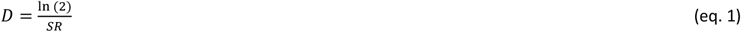

Where *D* is the population doubling time and *SR* the specific growth rate.

## 3. Results

### 3.1 Counting single *E. coli* cells using YOLOv4

We developed a method to characterize growth and lysis of bacterial populations by counting single bacterial cells using label-free bright-field microscopy and YOLOv4 based object detection oriented deep learning. Lysis of bacterial cells was induced upon infection of bacterial populations with their virus called bacteriophage or phage. In our experimental system, we employed the most studied bacterium, *Escherichia coli* to investigate bacterial growth in droplets, or infected *E. coli* with their phage T7 to observe bacterial cell death due to lysis. We employed the *E. coli* strain BW25113, which is a derivative of the wild-type *E. coli* K-12 strain, described as weakly motile, and it is a known host strain for propagation of phage T7[46, 47]. T7 is the most suitable phage model for bacterial real-time lysis experiments because of its short infection cycle (approx. 15 minutes at 37°C) and high lytic activity [48]. Using our method, individual *E. coli* cells were trapped in anchored droplets and their growth and lysis observed in a 3D liquid environment. The YOLOv4 model to detect cells in focus was trained for 6000 iterations and the average precision (AP) value of 83.6% was obtained (Figures S3 and S4). For each experiment, one anchored droplet was selected and observed over a period of up to 10 hours. Droplets were imaged using a Z-stack method. Briefly, 20 to 30 images at 0.5 microns intervals were acquired, ensuring that cells were in focus and detected by our model in at least one focal plane in the stack (c.f. section 2.4). The whole stack was acquired in 1.5 to 2.1 seconds depending on the number of slices chosen. Z-stack images were circularly masked to avoid any detections outside the zone of interest (see Supplementary section 8). The masked images were used as the input for the trained YOLOv4 model and detection images were obtained along with coordinates of all bounding boxes. YOLO detections for each Z-stack generated the detection counts which were then processed using the 3^rd^ maximum or 3^rd^ minimum method to identify individual cell doubling and cell lysis events, respectively (see section 2.4 and Figure S9).

Figure 3 shows representative bacterial growth and bacteriolysis assays. Figure 3A shows the increase in cell count over time. Each step in the processed count plot represents cell division events. Increase in cell count for each step was manually verified using corresponding time-point images as seen in Figure 3B and Supplementary Movie S1. Similarly, Figure 3C shows a decrease in cell count as phages cause bacteriolysis. Each step corresponds to lysis events and was verified using corresponding time-point images as seen in Figure 3D and Supplementary Movie S2.

### 3.2 Analysis of *E. coli* growth

Experiments were conducted at different temperatures and different initial cell numbers to check the versatility and applicability of the cell count method developed using the YOLOv4 model. For all experiments, we visualised *E. coli* cells growing in culture medium in the 3-dimensional environment of the anchored droplet. Table 1 gives a detailed account of experimental conditions employed during growth experiments. The parameters for imaging of each of these growth experiments have also been listed in Table S1. All growth curves include a 10-minute delay due to the time required for cell loading and initiation of image acquisition.

#### 3.2.1 Growth of *E. coli* at room temperature

We conducted multiple experiments at room temperature starting at various loading optical densities corresponding to different bacterial population size and initial cell numbers. Mean cell number could be anticipated from Poisson statistics using a droplet volume of 15 pL calculated assuming an ellipsoid droplet shape. In the first experiment, we started with a loading OD_600_ value of 0.19 and an initial cell count of 2. We reached a final cell count of 47 after 470 minutes. When the loading OD_600_ was 0.35 and initial cell count was 1, cell number reached a final count of 37 in 396 minutes. When we started with a loading OD_600_ of 1 and an initial cell count of 5, we reached a final count of 21 in 445 minutes. Figure 4A shows the increment in cell count in the form of individual steps every time a cell divides for each of these experiments. We observed that the growth of cells in each condition was dependent on the loading OD_600_ values in all the cases. Single-cell times to division in all experiments were broadly evenly spread across the duration of the experiments as seen in Figure 4B. Population doubling times of 103 ± 3 minutes (OD_600_=0.35), 156 ± 3 minutes (OD_600_=0.19) and 215 ± 5 minutes (OD_600_=1) were calculated. The growth of cells in the droplets was found to be reflective of the media carrying capacity. At lower OD_600_ values (0.19 and 0.35), we observed faster cell division and higher cell yield due to higher concentration of nutrients present, whereas droplets generated with high initial OD_600_ showed slower growth and lower yields due to nutrient depletion in the media as media carrying capacity was approached.

**Figure 4.**
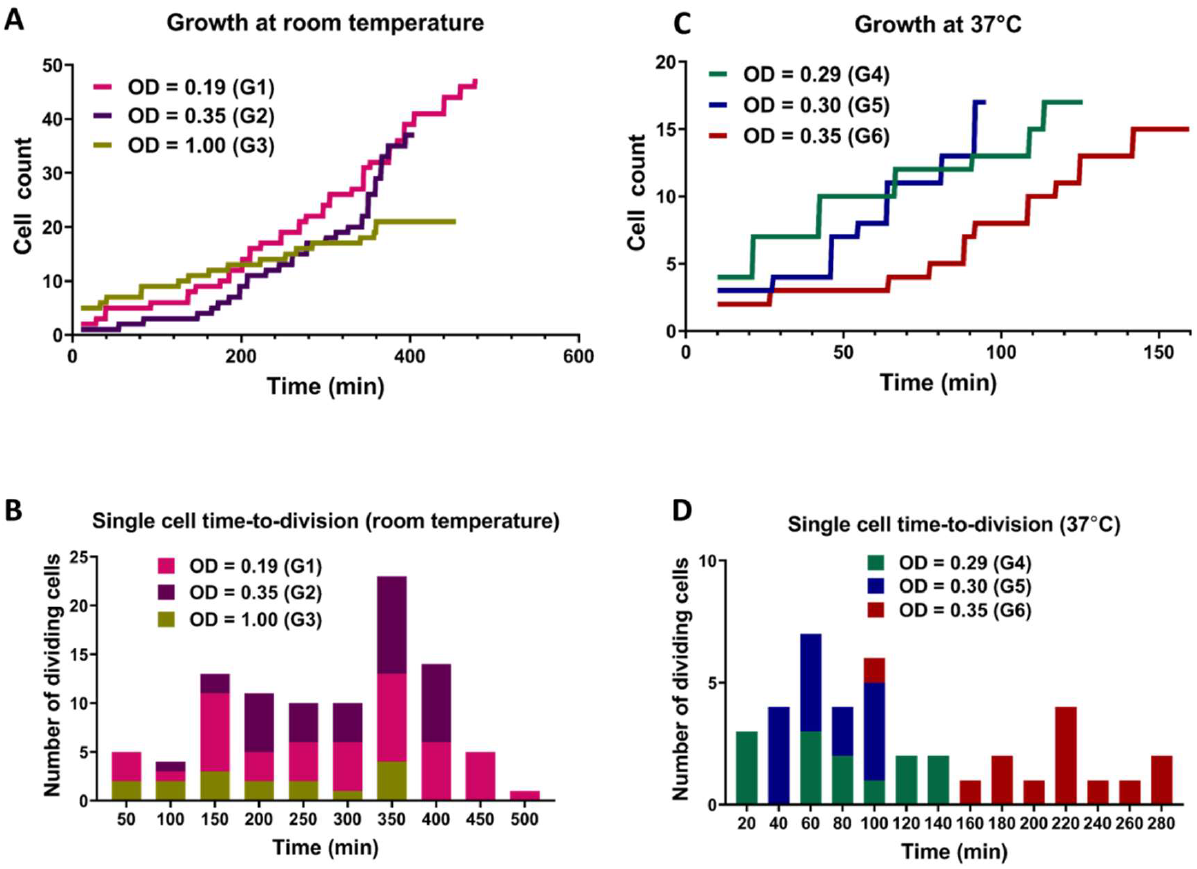
Comparison between growth of *E. coli* cells at room temperature and 37 °C. **(A)** Growth of *E. coli* populations at room temperature determined by the increase in number of cells due to cell division. **(B)** The corresponding histogram showing the distribution of single-cell time-to-division for each experiment. **(C)** Growth of *E. coli* populations at 37 °C determined by increase in number of cells due to cell division. **(D)** The corresponding histogram showing the distribution of single-cell time-to-division.

#### 3.2.2 Growth of *E. coli* at 37°C

We performed three experiments starting with different OD_600_ and number of bacterial cells in droplets. When the loading OD_600_ were 0.29 and 0.30 with cell count of 4 and 3 respectively, the cell population reached the same count of 16 in 135 minutes and 100 minutes respectively. When the OD_600_ was 0.35, and the droplet contained 2 cells initially, the cell number reached a total of 15 in 145 minutes. Since the media carrying capacity was almost identical in all three cases, the growth yields for all experiments were comparable. Figure 4C shows the increment in cell count in form of individual steps for experiments conducted at 37°C. The doubling times were calculated to be 29 ± 3 minutes (OD_600_=0.30), 43 ± 3 minutes (OD_600_=0.35) and 79 ± 5minutes (at OD_600_=0.29).

### 3.3 Analysis of *E. coli* lysis with T7 phage

Phage T7 was chosen to perform bacteriolysis experiments as its lytic cycle causes the infected cells to undergo sudden burst, making identification of intact cells that retained their morphology easier.

#### 3.3.1 Lysis of *E. coli* cells at room temperature

Aliquots of exponentially growing *E. coli* cultures with determined OD_600_ values were mixed with the T7 phage lysate (10^9^ PFU/mL) at different volumes to obtain different expected MOI in droplets for each experiment. Specific parameters of all lysis experiments are summarized in Table 2. The initial cell counts in experiment with expected MOI value 0.35 was 28 and all the cells in the observed droplet were lyzed within 206 minutes. In the experiment with an expected MOI value of 0.45, we started the experiment with an initial cell count of 29 and noted complete bacteriolysis in 123 minutes. For the droplets with expected MOI value of 0.48, complete cell lysis from an initial cell counts of 17 was observed within 106 minutes. Figure 5A shows the comparison for the three lysis experiments at room temperature. We observed that higher MOI resulted in faster cell lysis, which is to be expected as applying higher MOI indicates that bacterial cells were initially challenged with more phage particles. Figure 5B shows individual cell lysis events extracted from Figure 5A. A peak of lysis at time 60 minutes seem to point towards the average time to lysis of *E. coli* at room temperature. Indeed, in experiments L2 and L3, we observed a sudden onset of cell burst at 60 minutes, lyzing more than half the population of cells.

**Figure 5.**
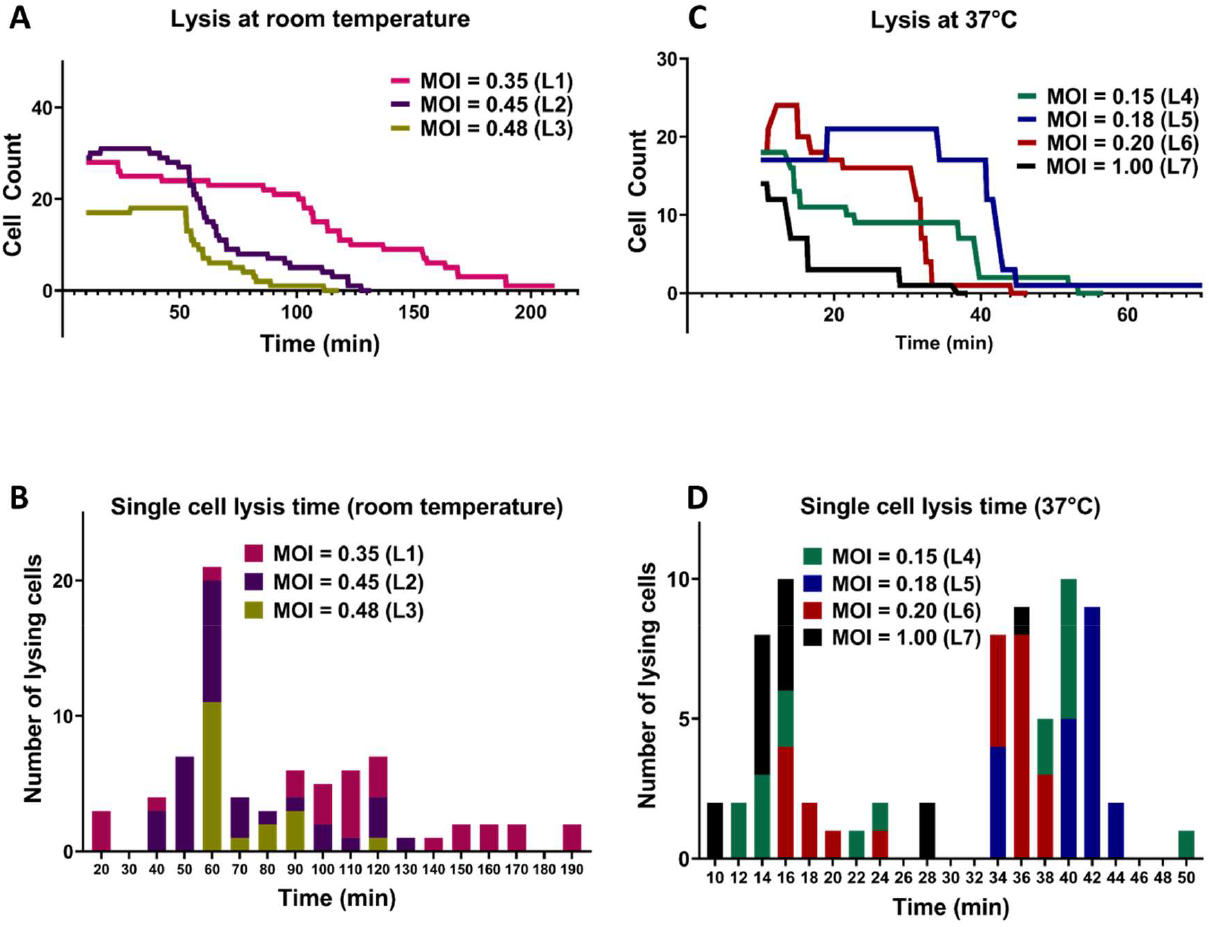
Comparison between lysis of *E. coli* cells by T7 phages at room temperature and 37 °C. **(A)** Lysis of individual *E. coli* cells at room temperature. **(B)** The corresponding histogram shows the distribution of time to lysis of single bacterial cells for each experiment. **(C)** Lysis of individual *E. coli* cells at 37 °C. **(D)** The corresponding histogram shows the distribution of time to lysis of single bacterial cells. All lysis curves include a 10-minute delay due to the time required for cell loading and initiation of image acquisition.

#### 3.3.2 Lysis of *E. coli* by T7 phages at 37°C

The lytic cycle of the T7 phage at 37°C is known to be around 17 minutes resulting in the duration of lysis experiments at 37°C being significantly shorter than that of lysis experiments at room temperature [49, 50]. Considering the speed of T7 lytic cycle, low expected MOI values were chosen to aid in observing individual lysis events. Parameters of experiments and imaging are summarized in Table S2. For droplets with MOI value 0.15, we began the experiment with an initial cell count of 18 (i.e. estimated initial phage count of 3) and observed complete cell lysis in 48 minutes. For droplets with expected MOI value 0.18, the initial cell count was noted to be 17 (estimated initially as 3 phage particles) and almost all the cells were lyzed within 39 minutes. The T7 phage burst size refers to the number of viral particles produced during the lytic cycle of phage infection and is predicted to be hundreds for T7 phage[51]. However, one cell was observed to remain whole even with the expected presence of large number of T7 phages produced by previous lysis of other cells in the same droplet (experiment L5).

For droplets with expected MOI 0.20 with initial cell count 18, cell count increased to 22 (experiment L6) and most cells lyzed after around 30 minutes with complete lysis observed within 38 minutes. For the experiment with a relatively high MOI of 1 (experiment L7) and initial cell count 14 (14 expected phages initially), approximately 75% of the population was lysed within 16 minutes and complete cell lysis was observed in 36 minutes. Figure 5C shows the comparison between lysis experiments at different expected MOI. Similar to room temperature experiments, we observed faster cell lysis in droplets with higher expected MOI. Figure 5D shows individual cell lysis events for each experiment in Figure 5C. Overall, we observed both early lysis (between 11 and 17 minutes) and late events with noticeable time delay as seen in Figure 5D (with many recorded lysis events occurring after 28 minutes from mixing).

## Discussion

By combining bright field microscopy, microfluidics, Z-stack imaging and YOLOv4 based object detection framework, we demonstrate a robust platform that enables single-cell level monitoring of growth or lysis of bacterial cells in liquid environments. Growth experiments demonstrated the ability of anchored droplets to act as independent microreactors allowing the cells to grow with little physical constraints. This label-free platform enabled us to detect and count individual cells purely based on their morphological features without any biological modifications or need for fluorescent labelling. Our method poses limited restrictions (presence of an oil-water interface, finite resources within the droplet) on the cells and allows us to monitor large cell populations which more faithfully represent both *in vivo* and environmental conditions.

A key enabling advance has been the ability to detect cells of various morphology using deep learning methods. Our YOLOv4 model exhibited adaptable and accurate detection of *E. coli* cells in focus across different levels of brightness and contrast (Figure S6) making the method robust to experimental drift. Average Precision (AP) was calculated to be 83.6% with an average loss of 0.4 after 5600 iterations of training (see Figure S3 for more details). The weights corresponding to these conditions were used to perform all deep learning-based detections and analysis throughout this study. The method we developed using the coordinates of the cell detections across the Z-stack for each droplet aided in the analysis of individual cell positioning, distribution pattern and counts in a 3D environment.

The accuracy of detections was mainly limited by the positioning of cells with respect to the droplet interface and by the local density. Supplementary sections 8 and 9 exemplify false positive detections and show standard deviation as a function of cell number. We observed that cells were often undetected if they were positioned or moved around the circumference of the selected droplet as they could not be resolved in images as seen in image in Figure S7. Similarly, the accuracy in cell count was found to be directly dependent on the number of cells. Figure S8 shows the standard deviation for cell count accuracy versus the total number of cells. We observed that the accuracy of cell count decreased as the number of cells in the droplet increased, imposing a practical limit on the maximum number of cells that can be counted per experiment. Our radius of exclusion method also set a hard boundary on the maximum cell density (i.e., one cell per 44 µm^2^) corresponding to about 50 cells given our droplet diameter.

T7 phages cause infected *E. coli* cells to burst. The YOLOv4 model did not detect the lyzed cells or floating debris because it was specifically trained to detect cells that retain their shape as seen in Figure S2. One limitation of our method is the finite tracking ability of cells on an individual basis to construct the lineage tree of a population. This is linked to the relatively long time required to acquire complete Z-stacks, limiting our time resolution to typically 1.5-2.1 seconds. The use of smaller Z-stacks and shallower traps could increase this time resolution further and enable tracking of low-motility strains. Extension of the presented method to larger populations in larger drops (i.e., with larger, taller traps) could be implemented by adapting the method to evaluate cell-to-cell distances in 3D instead of the exclusion radius method presented. Automated screens of multiple droplets at a time could also be done using a motorized microscope stage at the cost of time resolution per droplet. Bacterial growth starting from a few cells exhibited strong stochasticity. As the histograms in Figure 4B illustrate, population development did not follow reproducible patterns, in line with expected stochasticity at the single-cell level which has been covered in detail in other studies [17, 21, 23]. Experiments at both room temperature and 37°C show stochasticity under similar growth conditions.

For phage experiments, we hypothesize that all cells in a given droplet may not be infected immediately especially at low MOI experiments. This allows non-infected cells to grow and divide as seen in Figure 5A and 5C (experiments L2, L5 and L6). As infected cells lyze, the concentration of phages in the droplet increases, causing the rest of the cells within the droplet to lyze from secondary infections as seen in Figure 5C.Phage life cycle highly depends on temperature. At room temperature, as found in the environment, phage adsorption and replication dynamics may be affected. Our experiment at room temperature still shows efficient lysis albeit much slower than at 37 °C. In our droplet microfluidics setup, T7 phage lyzed almost all cells within 200 and 50 minutes at room temperature and 37 degrees respectively. However, some cells did not lyze for several minutes even after the lysis of all other cells and remained motile in the droplet (Figure 5C, experiment L5). This could be explained by resistance against phage infection developed via many different mechanisms [52].

## Conclusions

Our microdroplet trapping device and analysis pipeline from Z-stack bright-field images provides a means to study dynamic interactions between microbes and antimicrobial agents. This includes the life cycle of phages, e.g., dynamic transition between lysogenic and lytic pathways observed via long-term balance between growth and lysis, and the evolution of resistance to phages, an essential consideration for pre-empting future resistance to phage treatment. The technical simplicity of the setup coupled with shareable deep learning models will accelerate phenotypic studies of bacterial populations. In future work, the platform could be used to study combinatorial effects of antibiotics and phages. One could also expand object detection-oriented methods to study polymicrobial cultures of clinically relevant strains to characterise the impact of a microbial community composition with respect to the lytic efficacy of phages. Finally, our method could be used to evaluate susceptibility of cells to antimicrobial agents other than phages via change in bacterial growth dynamics or morphological changes at sub-inhibitory concentrations.

## Acknowledgements

The authors thank Dr. Remy Chait for useful discussions and comments, Dr. Wolfram Möbius for the bacterial strain and phage. We would also like to acknowledge the use of Exeter Microfluidics Facility.

## Data availability statement

Code used for YOLOv4 training and conversion of Darknet weights to TensorFlow are available on https://github.com/AlexeyAB/darknet

https://github.com/theAIGuysCode/yolov4-custom-functions

All the MATLAB scripts used for analysis in this paper and Python code for Z-stack acquisition and drift correction are available at:

https://github.com/GielenLab/YOLOv4-bacteria-analysis

Training images and model weights data will be shared upon reasonable request.

