## Supplementary Information for "Label-free analysis of bacterial growth and lysis at the single-cell level using droplet microfluidics and object detection-oriented deep learning"

### 1. Imaging parameters for each experiment

Tables S1 and S2 give information on the imaging conditions for all the growth and lysis experiments. As seen in the tables, the average time taken to acquire individual Z-stacks varied between 1.5 second and 2.1 seconds depending on the total number of images in the stack. As bacterial growth is slower at room temperature, the time gap between stacks at room temperature was chosen to be higher than for experiments at 37°C. We followed the same pattern for lysis experiments as lysis at room temperature took significantly longer than lysis at 37°C.

| Experiment | Number of slices in Z-stack | Distance between each slice (µm) | Time to acquire one stack (s) | Total height of Z-stack (µm) | Time gap between each Z-stack (s) |
|---|---|---|---|---|---|
| G1 | 20 | 0.5 | 1.5 | 10 | 60 |
| G2 | 25 | 0.5 | 1.9 | 12.5 | 60 |
| G3 | 24 | 0.5 | 1.8 | 12 | 60 |
| G4 | 28 | 0.5 | 2.1 | 14 | 10 |
| G5 | 28 | 0.5 | 2.1 | 14 | 10 |
| G6 | 28 | 0.5 | 2.1 | 14 | 10 |

**Table S1: Imaging parameters for growth experiments**

| Experiment | Number of slices in Z-stack | Distance between each slice (µm) | Time to acquire one stack (s) | Total height of Z-stack (µm) | Time gap between each z-stack (s) |
|---|---|---|---|---|---|
| L1 | 28 | 0.5 | 2.1 | 14 | 60 |
| L2 | 28 | 0.5 | 2.1 | 14 | 60 |
| L3 | 28 | 0.5 | 2.1 | 14 | 60 |
| L4 | 28 | 0.5 | 2.1 | 14 | 10 |
| L5 | 28 | 0.5 | 2.1 | 14 | 10 |
| L6 | 28 | 0.5 | 2.1 | 14 | 10 |
| L7 | 28 | 0.5 | 2.1 | 14 | 10 |

**Table S2: Imaging parameters for lysis experiments**

### 2. CFU/PFU assay

##### 2.1 Colony Forming Unit assay (CFU)

Aliquots from exponential culture used for droplet experiments were obtained and serially diluted in the LB media (10 g/L tryptone, 5 g/L yeast extract, 10 g/L NaCl) by factors from $10^{-1}$ to $10^{-8}$ in a microwell plate. 100 µL of dilutions $10^{-6}$ and $10^{-7}$ were plated on LB agar plates and left overnight at 37° C. Bacterial colonies were counted the next morning to obtain population size (CFU/mL) used for the droplet experiments.

##### 2.2 Plaque Forming Unit assay (PFU)

Soft agar is primarily used in a laboratory technique called the soft agar assay or colony formation assay. In the soft agar assay, cells of interest are mixed with the liquid agarose solution and then plated onto a solid agarose layer in a petri dish. The dish is incubated, allowing the cells to proliferate and form colonies on the surface of soft agar. The soft agar layer prevents cell migration and forces the cells to grow in a restricted manner, which aids in the visualization and quantification of colony formation. An exponential culture of *E. coli* BW25113 was prepared in LB medium supplemented with 2 mM $CaCl_2$ by diluting a previously overnight culture at a 1:1000 ratio. The culture was then grown for approximately 5.5 hours to allow for optimal growth. To prepare phage lysate dilutions, a 96-well plate was used. In a 15 mL Falcon tube, 20 µL of GLC glucose-$CaCl_2$ mix (1 M glucose and 2.5 mM $CaCl_2$ in final concentrations) was added, followed by the addition of 200 µL of the *E. coli* culture. Subsequently, 4 mL of soft agar was introduced to the Falcon tube. The mixture was immediately vortexed briefly to ensure proper mixing and poured onto an agar plate. After pouring, the plate was left at room temperature for approximately 2-5 minutes to allow the agar to solidify. Once solidified, 5 µL of the phage dilutions of factors $10^{-1}$ to $10^{-11}$ were spotted onto the plate. The plate was then left on the bench for 1-2 hours to further solidify and left at room temperature overnight for incubation. Plaques were counted the next morning to calculate the concentration of phage titer in PFU/mL.

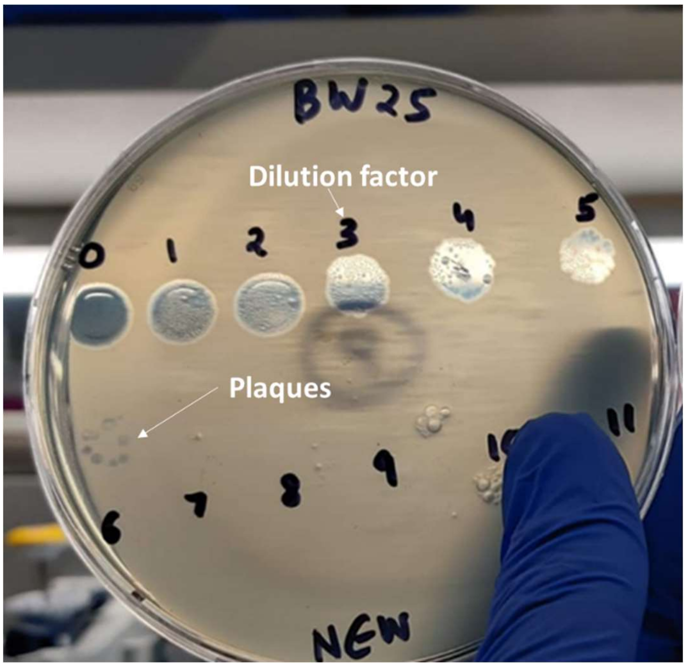

**Figure S1:** PFU assay to calculate concentration of T7 phage titer.

#### 3. Labelling of training set for supervised learning

Each cell of interest was labelled as 'in-focus' cell based on the morphology as seen in the Figure S2 (a), (b), (c) and (d). Cells were in focus if a clear outline was visible along with the characteristic rod shape. Images were pooled from multiple experiments that have been performed on different days and with different imaging conditions (e.g., brightness, contrast) to generate robust models.

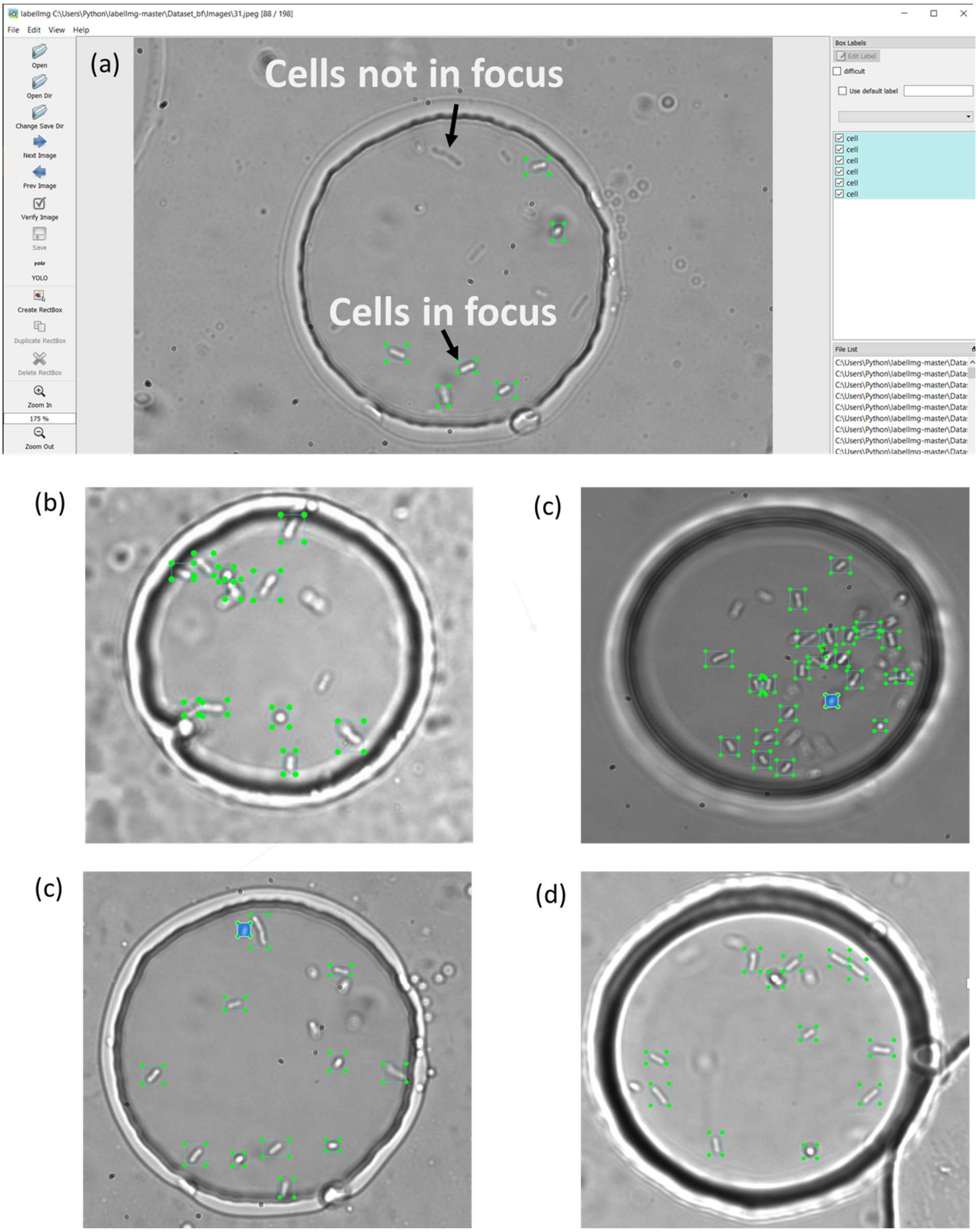

s

**Figure S2:** Examples of labelling of cells in focus using LabelImg. Bounding boxes are manually drawn around each cell.

#### 4. YOLOv4 model selection

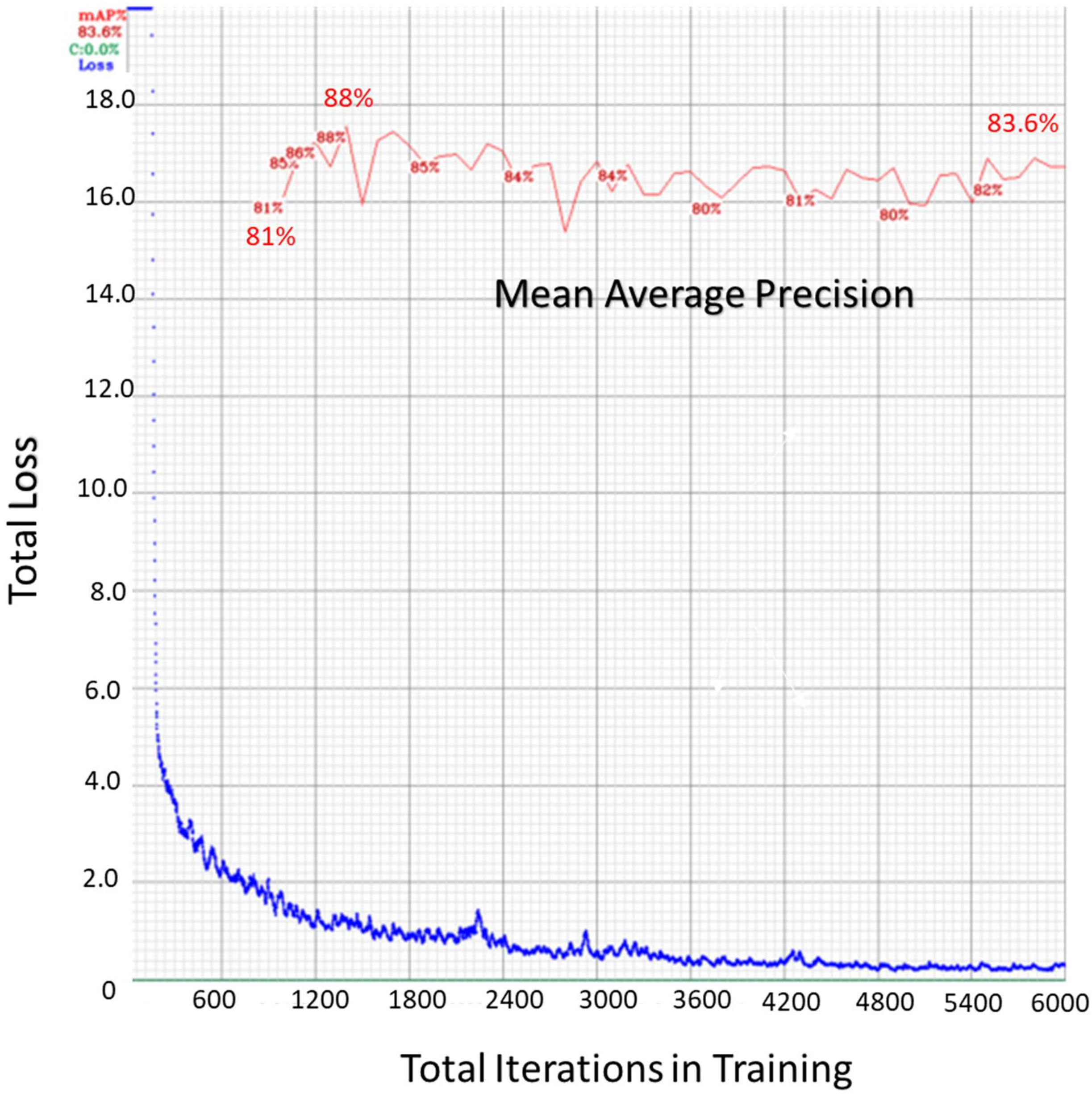

**Figure S3:** Yolov4 training curve for a single object of interest (cell) based on manually labelled training data as shown in Figure S3. Final weights were selected at 5600 iterations.

YOLOv4 training was performed on the Google Colab platform. The dataset consisted of 200 images with over 1670 single cells labelled. The total labelling and training time were 5 hours and 8 hours, respectively. The weights of the training were backed up every 100 iterations. An epoch in machine learning is defined as one complete run-through of the entire dataset by the algorithm. The best mAP value was obtained after 1500 iterations (~50 epochs) of training at 88.1%. However, the average loss at this point is significantly higher than the desired value of less than 0.5. Therefore, we chose the weights once the loss was sufficiently low, after 5600 iterations (~187 epochs). The total loss used for the final model in all experiments after completion of the training was calculated to be 0.301 with a corresponding mAP of 83.6%.

#### 5. Precision, recall and IOU

In YOLOv4, precision and recall are calculated using the standard definitions from the field of object detection. Precision is the fraction of true positive detections out of all positive detections made by the model. Mathematically, it can be expressed as:

**Precision = True Positive / (True Positive + False Positive)**

where True Positive is the number of correctly predicted objects, and False Positive is the number of objects predicted by the model that do not exist in the ground truth data.

Recall is the fraction of true positive detections out of all ground truth positives. Mathematically, it can be expressed as:

**Recall = True Positive / (True Positive + False Negative)**

where True Positive is the number of correctly predicted objects, and False Negative is the number of objects in the ground truth data that were not detected by the model.

The F1 score is typically used to evaluate the accuracy of object detection models by comparing the predicted bounding boxes to the ground truth bounding boxes. Higher F1 scores indicate that the model is better at accurately detecting objects in images, while lower F1 scores indicate lower accuracy. Yolov4 can be trained using various techniques such as data augmentation, transfer learning, and more, to improve the F1 score of the model.

**IOU (Intersection over Union)** is a metric used to evaluate the accuracy of object detection models, including YOLOv4. It measures the overlap between the predicted bounding box and the ground truth bounding box for an object in an image. In YOLOv4, the IOU is defined as the ratio of the intersection area between the predicted bounding box and the ground truth bounding box to the union area of these two boxes. It is expressed mathematically as:

**IOU = (Area of intersection) / (Area of union)**

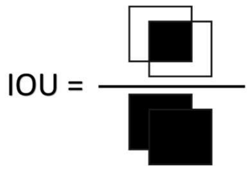

An IOU value of 1 indicates that the predicted bounding box perfectly overlaps with the ground truth bounding box, while a value of 0 indicates that there is no overlap. In YOLOv4, a threshold IOU value is used to determine whether a detection is considered a true positive or a false positive. If the IOU is greater than the threshold value, the detection is considered a true positive, otherwise it is considered a false positive. Due to the size of the images and cells within these images, significant IOU variation is caused by very few pixels. In our case, the IOU threshold of 50% with 83.6% detection accuracy was found satisfactory in detections of cells.

The mean average precision (mAP) or average precision (AP) in our case (since we have only one class of objects being detected) was measured at different IOU percentages demonstrating the effectiveness of the model. Figure S4 shows the comparison of between different model parameters used to evaluate the performance of the trained YOLOv4 model.

(a)

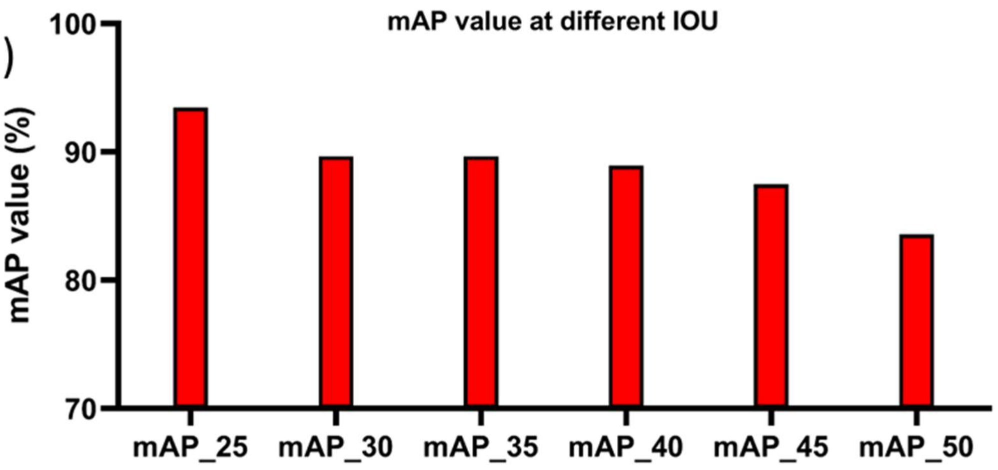

(b)

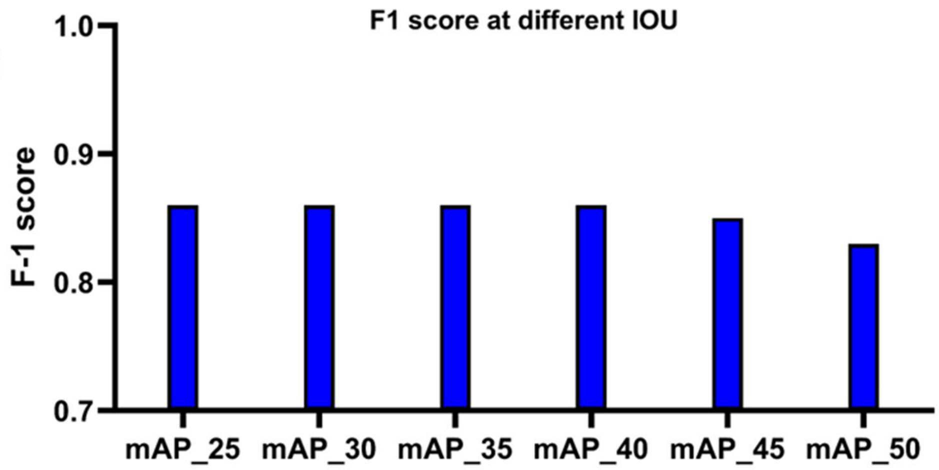

**Figure S4**: mAP and F1 score at different IOU. The mAP_X labelled in x-axis of the plots in Figure S4 corresponds to X% IOU in the model. (a) mAP at different IOU thresholds for the model selected for experimental analysis (b) F1 score curve for the final weights of the YOLOv4 model. Values over 0.8 for mAP reflect satisfactory model performance.

Figure S4(a) gives a comparison between the accuracy of detection of the model weight selected for experimental analysis at different IOU thresholds. The average precision of the model is 83.6% even at 50% IOU. The F-1 score is shown in Figure S4(b) is the harmonic mean of precision and recall. Higher F-1 score reflect better model performance and an F-1 score of >0.80 was obtained at an IOU threshold of 50%.

#### 6. Detections across the Z-stack

Figure S5 shows the overlay of a single Z-stack showing the positions and number of detections of each individual cell in the stack. The green dot represents the centroids of the detection box of individual detections in each plane in the z stack. The red dot represents the final position of that particular cell calculated based on the positioning of individual green dots within an arbitrary radius. This arbitrary radius is called radius of exclusion.

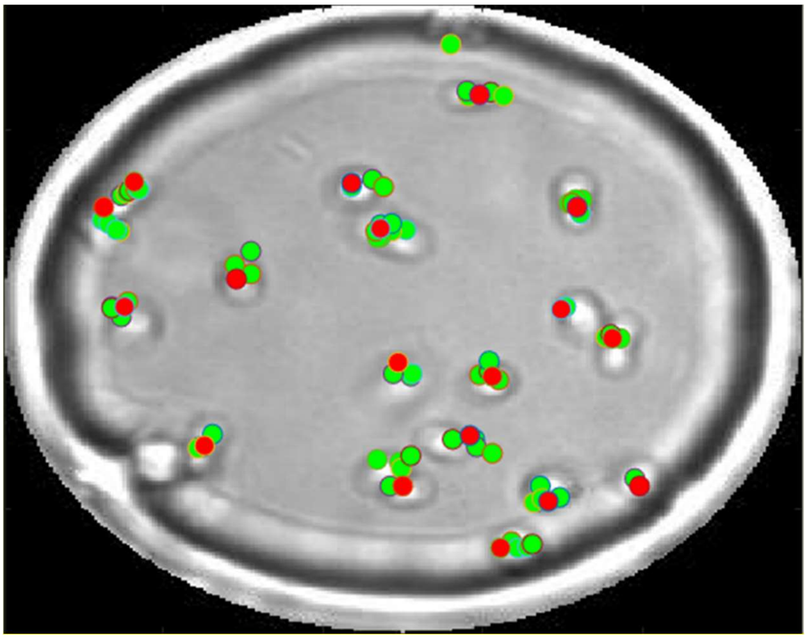

**Figure S5**: Z-stack overlay for a single time point after implementing the counting method.

#### 7. Detections under varying imaging conditions

Light intensity to image all droplets is kept constant but it is often difficult to replicate imaging conditions of droplets due to their positioning, placement of the condenser as well as the positioning of the main focal plane. Therefore, the YOLOv4 model needs to be robust to perform accurate detections even under different imaging conditions such as varied contrast, brightness and pixel intensity distributions (including pixel saturation). Figure S6 given examples of accurate detections under varied brightness and pixel intensity distributions showing that our model performs well even with different imaging conditions.

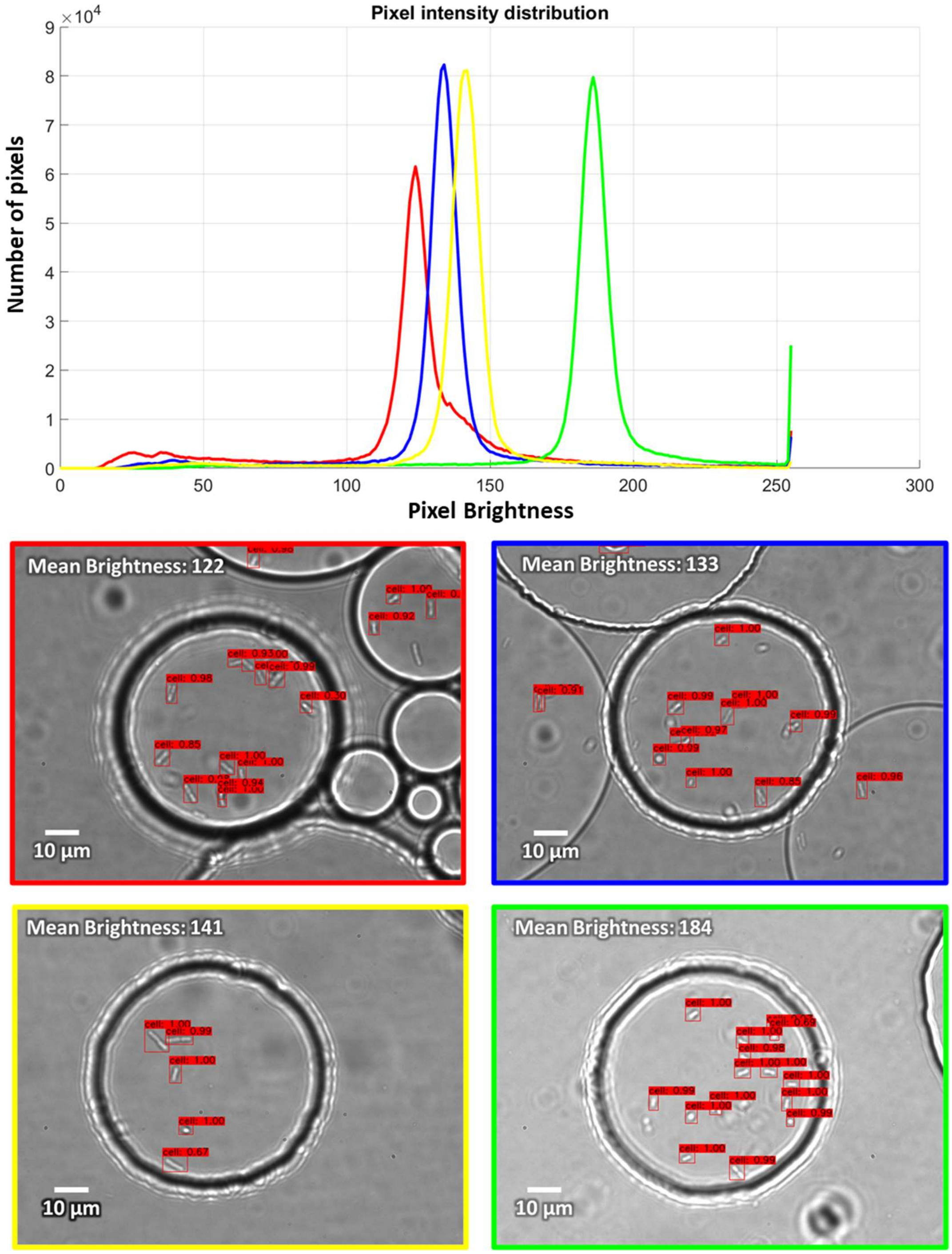

**Figure S6:** Examples of cell detections with confidence values. (a) Pixel intensity distribution and (b) corresponding images with average brightness values. Images were part of the experimental dataset (not part of the training set) demonstrating the robustness of the YOLOv4 model.

#### 8. False detections in the masked droplet

During data analysis, it is important to focus on a single droplet. To this end, we applied a mask to only analyse a single circular region of interest as shown in Figure S7. In some cases, we observed detections outside the droplet as seen in Figure S7(B). All the images obtained during experiments were masked to avoid any impact on the cell count from the detections outside the region of interest.

We also observe that some cells were not detected because of they are positioned close to the interface of the droplet and the trap (Figure S7A and S7B) or because they formed aggregates.

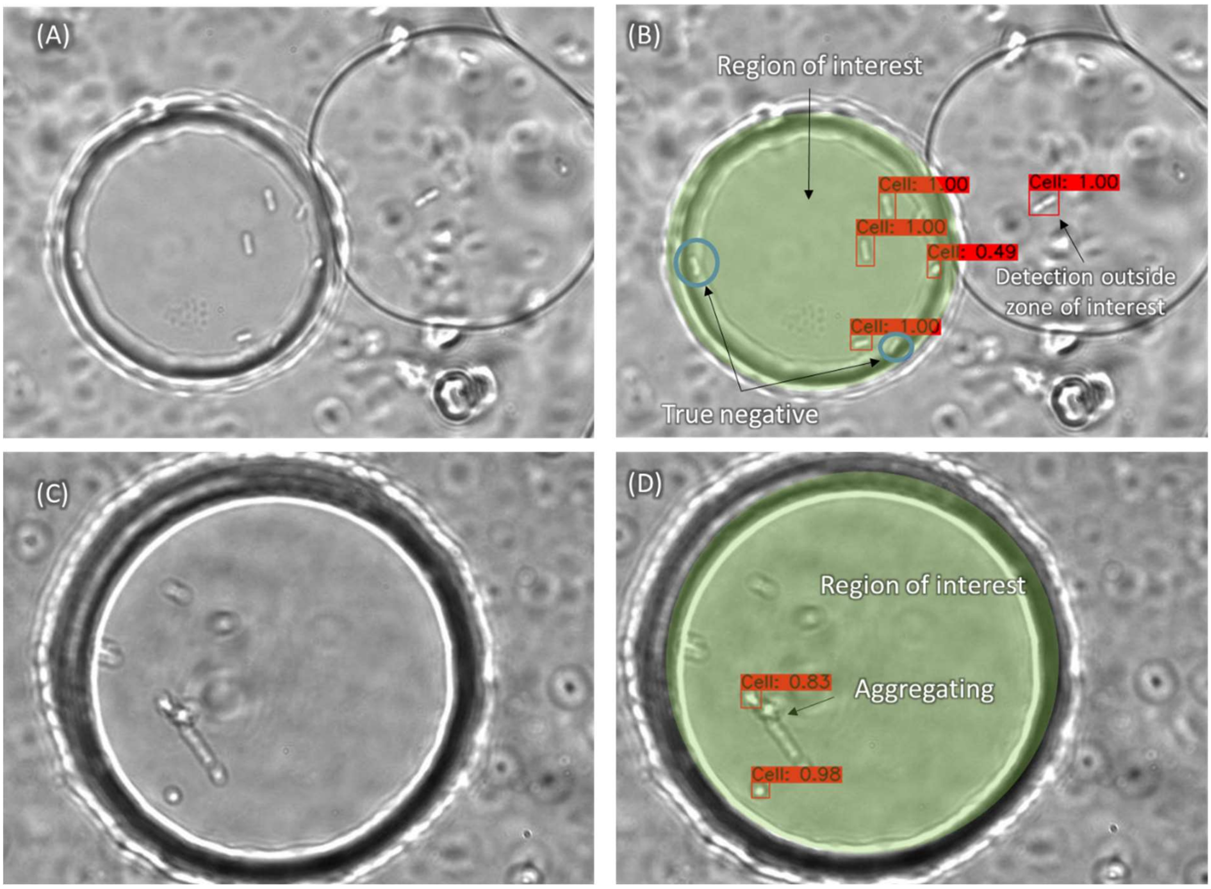

**Figure S7**: Examples of aggregates and detections outside zone of interest. (A) Raw image from an experiment before performing detections. (B) Image A after performing detections. We observe that some cells located near the droplet-trap interface are not detected. (C) Image of a droplet from an experiment showing aggregating. (D) Image C after performing detections. We observe that not all the cells are accurately detected due to the cell aggregation.

#### 9. Detection errors as a function of total cell number in a droplet

We observed that the error in cell count is directly related to the total number of cells. As the number of cells in the trapping chamber increased, the error associated with their count also increased linearly as shown in Figure S8. In other words, the lower the number of cells, the more accurate the generated cell count.

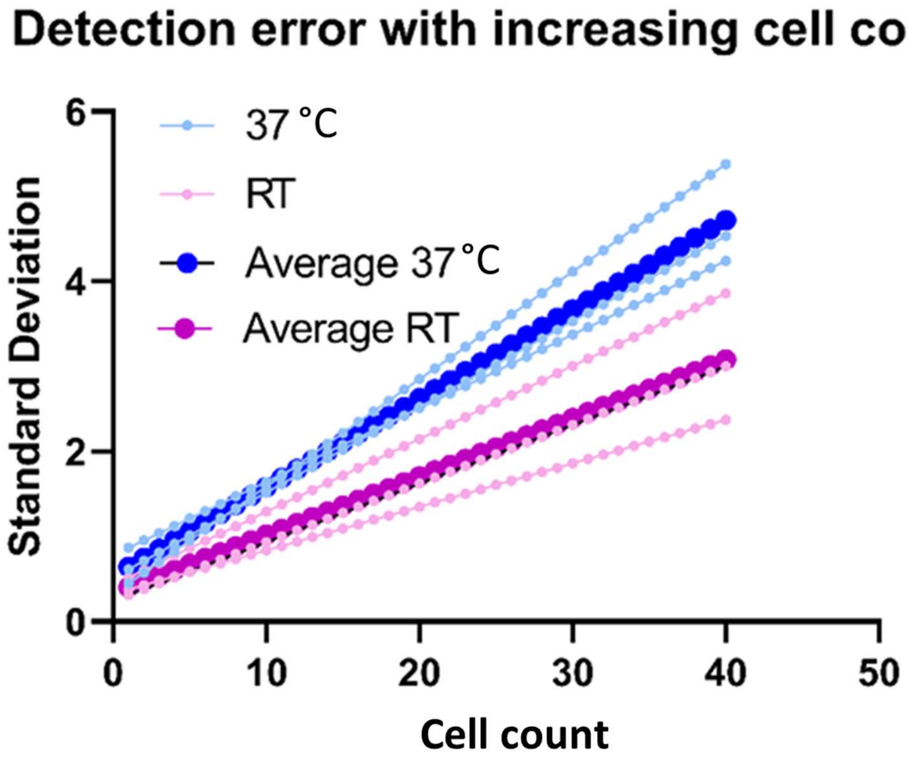

**Figure S8:** Standard deviation in cell number as a function of the total cell count at room temperature (RT) and 37 °C.

#### 10. Differences in single-cell time-to-division and time-to-lysis using the 1st to 5th maximum and minimum method.

The differences seen in single-cell time-to-division and time-to-lysis using the 1st to 5th maximum and minimum method are visualized in Figure S9. These methods prevented counts from being sensitive to local fluctuations due to false detections.

(a)

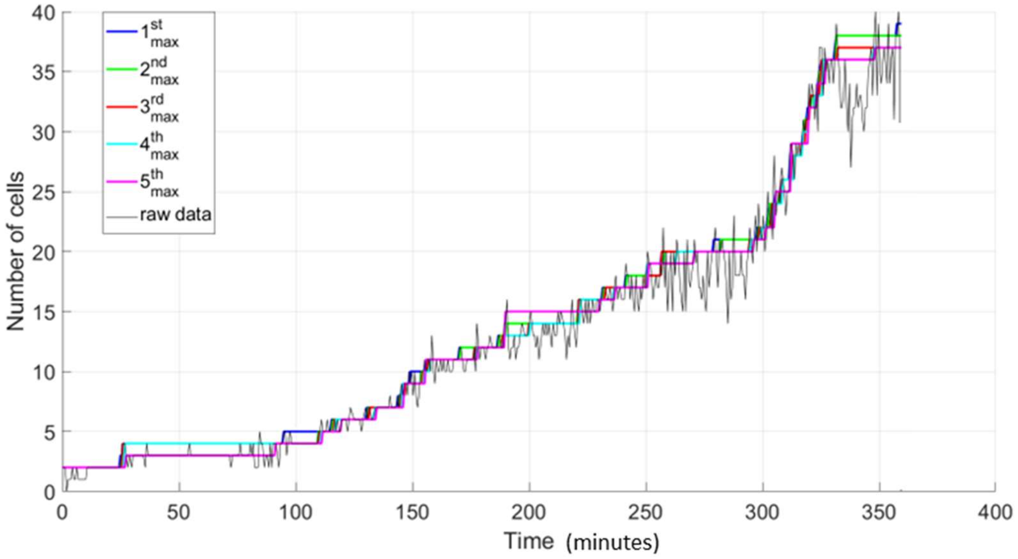

(b)

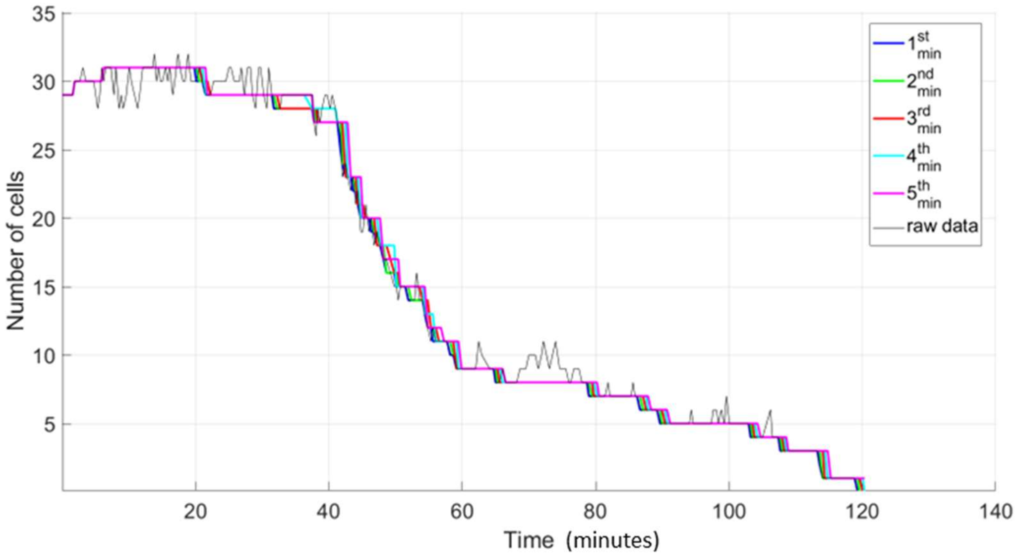

**Figure S9**: Comparison between detection count and processed count based on varying the maximum and minimum coefficient. (a) The processed counts for growth experiment 2 using the $n^{th}$ max method (n=1,2,3,4,5) (b) The processed counts for lysis experiment 2 using $n^{th}$ min method (n=1,2,3,4,5).

Based on these plots and comparison of images corresponding to each time point, we chose the $3^{rd}$ max (growth experiments) and $3^{rd}$ min (lysis experiments) to be the most accurate reflection of counts observed in the raw data. The curves corresponding to growth and lysis in Figures 4 and 5 were generated using the $3^{rd}$ max and $3^{rd}$ min processed counts.

#### 11.Calculation of doubling time for bacterial growth experiments

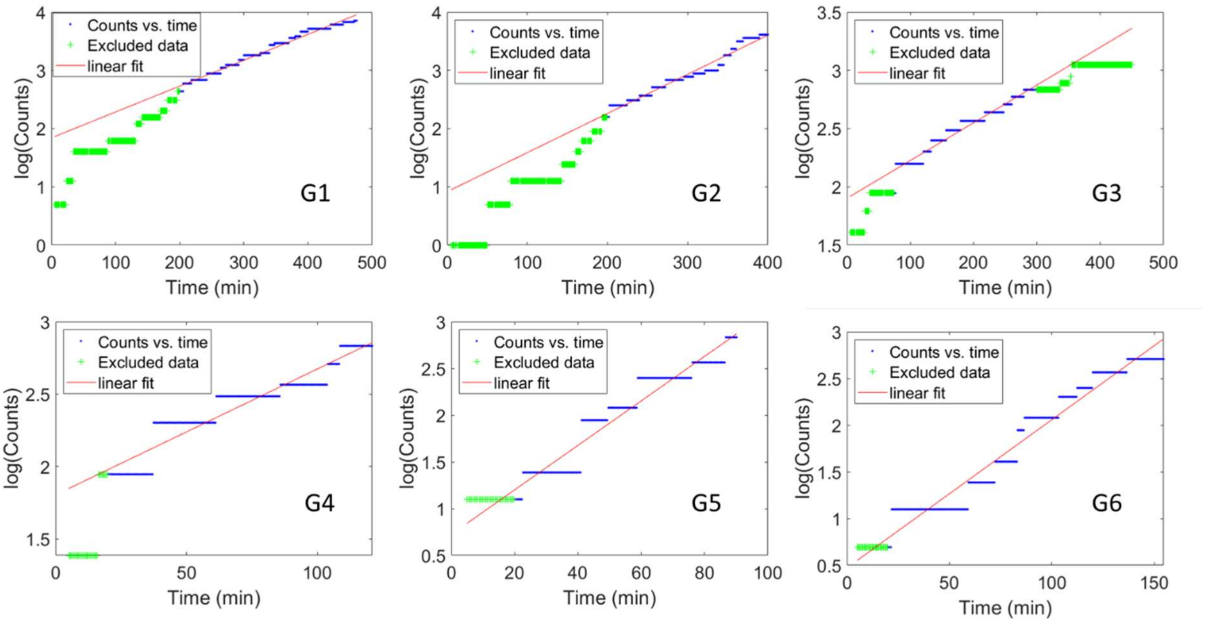

**Figure S10:** Calculation of doubling time from processed counts obtained for each growth experiment. Experiments G1, G2 and G3 were conducted at room temperature whereas experiments G4, G5 and G6 were conducted at 37°C.

#### 12. Movie captions

##### Movie S1

Growth of *E. coli* cells observed using our deep learning framework. Red dots represent the position of each cell and the increase in the number of red dots is reflective of an increase in cell count as seen in the movie. This movie is representative of a sample growth experiment with an initial cell count of 1 and a final cell count of 37 in 360 minutes.

##### Movie S2

Lysis of *E. coli* cells observed using our deep learning framework. Red dots represent the position of each cell and the decrease in the number of red dots is reflective of the decrease in cell count. The movie is representative of lysis experiment L2 with an initial cell count of 29 and complete cell lysis in 121 minutes.

##### Movie S3

Growth of *E. coli* cells as observed using the detection boxes form our deep learning framework. The red dots in the Movies S1 and S2 were calculated using the average position of the center of each detection box and their overlapping positions as seen in Figure S6. Green dots are the centroids of each individual detection box, and the red dots are the final calculated positions. The Movie S3 corresponds to the same dataset as in Movie S1.
